# Female budgerigars prefer males with foraging skills that differ from their own

**DOI:** 10.1101/2024.12.02.626337

**Authors:** Yuqi Zou, Zitan Song, Jiani Chen, Yuehua Sun, Michael Griesser

**Affiliations:** Key Laboratory of Animal Ecology and Conservation Biology, Institute of Zoology, Chinese Academy of Sciences, Beijing, 100101, China; Department of Collective Behavior, Max Planck Institute of Animal Behavior, 78464, Konstanz, Germany; Department for the Ecology of Animal Societies, Max Planck Institute of Animal Behavior, Konstanz 78467, Germany; College of Innovation Ecology, Lanzhou University, Lanzhou, 730000, China; Department of Biology, University of Konstanz, 78464 Konstanz, Germany; Center for the Advanced Study of Collective Behavior, University of Konstanz, 78464 Konstanz, Germany

**Keywords:** foraging skills, skill-pool effect, mate choice, social preference, *Melopsittacus undulatus*

## Abstract

Foraging skills influence food intake and may therefore also play a role in mate choice decision. Previous empirical work has shown that individuals benefit from being in groups that include individuals with a variety of foraging skills as this increases foraging success. This idea, formalized in the skill-pool hypothesis, may extend to mate choice. Diverse foraging skills can expand the foraging niche of a pair and benefit offspring through enhanced parental provisioning, and exposure to a broader foraging skillset. To test this idea, we trained captive female and male budgerigars to solve one of two different novel foraging puzzle boxes. Then, females simultaneously observed two males that could solve either the same or the other box, and assessed female preferences in a binary mate choice apparatus. Females preferred males with foraging skills that differed from their own, independent of the skill type and the number of times males solved the foraging puzzle. These findings show that foraging skills can influence social preferences, including in a mate choice context, and support intraspecific diversity in foraging skills.

## INTRODUCTION

Choosing a mate is a critical decision because mate quality and compatibility do influence fitness (Andersson and Simmons 2006). Mate choice decisions can be based on direct benefits, such as choosing a mate living on a high-quality territory or one with the ability to provide parental care (good parent hypothesis; Hoelzer 1989) and/or indirect benefits, such as selecting a mate that will produce attractive offspring (Fisherian sexy son hypothesis; Weatherhead and Robertson 1979) or possesses good genes (review in Kokko et al. 2003). In species with biparental care, males usually contribute to feeding the young and they can provide food to their mates during incubation. Accordingly, females should choose males that are efficient foragers, increasing the likelihood of breed successfully and raise more offspring. For example, female red crossbills (*Loxia curvirostra*) prefer males with higher feeding rates (Snowberg and Benkman 2009), and female spur-throated grasshoppers (*Melanoplus sanguinipes*) prefer males with better foraging abilities (Belovsky et al. 1996). In siskins (*Carduelis spinus*), male with longer yellow wing stripe, which is preferred by females, also solve foraging problems more quickly (Mateos-Gonzalez et al. 2011). Thus, mate choice based on traits linked to foraging efficiency can evolve.

Foraging skills (e.g., hunting or food processing techniques) affect foraging efficiency in many species (Patterson et al. 2015; Schuppli et al. 2016). For example, hunting skills in humans (*Homo sapiens*) are more important than physical strength for enhancing hunting efficiency (Gurven et al. 2006). Similarly in birds, immature olivaceous cormorants (*Phalacrocorax olivaceus*) exhibit lower foraging efficiency compared to adults, likely due to less developed foraging skills (Morrison et al. 1978). Evidently, foraging skills are crucial for fitness, as they directly influence energy acquisition. For example, survival in cactus finches (*Geospiza conirostris*) depends more on the ability to learn novel foraging skills than on beak morphology, which determines the foraging niche (Grant and Grant 1989). Moreover, a mate’s foraging skill is important not only for providing food but also because parents often serve as critical role models for offspring to learn from. Foraging skills are typically acquired from parents or conspecifics through social learning (Kulahci et al. 2016; van Schaik 2010), although they can also be acquired through innovation (Griffin et al. 2014). In birds, young learn their foraging skills primarily from their parents via social learning (Slagsvold and Wiebe 2011), and this process can continue post-fledging, facilitating the acquisition of more elaborate foraging skills (Uomini et al. 2020).

Some studies have explored female mate choice based on male foraging behavior, showing that females prefer morphological traits that are associated with foraging behavior. For example, female can choose mates based on carotenoid-based ornaments, which reflect male foraging success (e.g., Senar and Escobar 2002). However, there is limited evidence on whether direct observations of foraging behavior influence mate choice decisions (reviewed in Boogert et al. 2011). A recent experiment on budgerigars *(Melopsittacus undulatus*) showed that learning a novel foraging skill increases a male’s attractiveness for females (Chen et al. 2019). Assortative mating refers to the non-random selection of partners based on traits, which can be either similar (positive assortative mating) or dissimilar (negative assortative mating or disassortative mating). Positive assortative mating has been widely documented in relation to morphological traits (e.g., size and coloration) and physiological traits (e.g., age), whereas disassortative mating is rather rare. (Jiang, Bolnick, & Kirkpatrick, 2013). Notably, disassortative mating based on foraging skills remains understudied, although it could broaden the foraging niche of pairs (Beauchamp 2000; Matich et al. 2011) and reduce within-pair food competition (Selander, 1966). An example of disassortative mating is observed in a cichlid fish (*Perissodus microlepis*), where individuals pair with mates that have opposite mouth-opening directions, reduce competition by targeting different prey morphs (Takahashi and Hori 2008). Moreover, in species with biparental care where offspring acquire foraging skill from their parents, disassortative mating can increase the foraging skillset of offspring, thereby increasing their survival probability (Gibbons et al. 2005; Maisonneuve et al. 2021). This idea underlies the skill-pool hypothesis, which predicts that individuals should forage together with conspecifics that display different foraging behaviors within a group, as a diverse foraging skillset enhances overall foraging success (Giraldeau 1984). Specifically, foraging together with individuals that vary in their foraging skills, could benefit all group members via a broader range of food items and possibly achieving a higher foraging success. Indeed, foraging skill diversity increases food intake in house sparrow groups (*Passer domesticus*) (Aljadeff et al. 2020), which promotes the emergence and maintenance of foraging skill diversity (Firth et al. 2015).

Here, we used an experiment with captive budgerigars to assess whether unpaired females prefer unpaired males that had been trained in the same or a different foraging skill (i.e., extracting food from puzzle boxes). Budgerigars are social, group-foraging parrots that form enduring pair bonds (Stamps et al. 1985). Pairs remain together throughout the year, with males feeding their mate during incubation, while both parents provide food for nestlings and fledglings (Fritz 1976; Wyndham 1980). This suggests a role for foraging skills in mate choice. In the present study, males and females were trained to solve foraging puzzle boxes, with females learning either the same or a different skill than the males. After observing males extracting food from puzzle boxes, we assessed female preference for males with either same or different foraging skills to test whether females preferred males with different foraging skills, as predicted by the skill-pool hypothesis.

## METHODS

### Housing

Wild-type adult budgerigars (N=18 females, N=36 males, all > 1 year old) were obtained from commercial suppliers. Females and males were sourced from different suppliers to ensure unfamiliarity between birds of the opposite sex. Individuals were marked with numbered black leg bands and housed in same-sex cages under a 15.5 : 8.5 h light : dark regimen. The individuals in the cages were visually but not acoustically isolated. Females were first considered for an experiment once they had a brown cere, i.e., a physiological indicator of breeding readiness in budgerigars (Baltz and Clark 1996). All birds had *ad libitum* access to food (mixed millet seed), vegetables (cabbage, oil-seed rape leaves, and cucumber), fresh water, a cuttlefish bone, and grit, all sourced from online shops.

### Procedures

Two weeks before the experiments, subjects were moved from the same-sex cages to individual cages, where they were provided with the same environment as described above. We randomly selected one female and two males for each of the 18 experimental units. The experiment lasted 15 to 20 days (mean ± SE: 15.7 ± 0.4 days), and encompassed three phases (Table.1): (i) foraging skill training phase, (ii) observation phase, and (iii) preference phase. Given that one focal female was sick during the experiment and could not finish it, data from this experimental trio were excluded from the analyses.

**Table 1.** Timeline of the different test phases and location of birds.

| Phase | Duration | Location | Events |
| --- | --- | --- | --- |
| Phase 0 | 1 day | Individual cages | Select one female and two males. |
| Training phase | 7~12 days | Individual cages | Females and males were trained to open one puzzle box type. |
| Observation phase | 4 days | Individual cages | Females observed males extracting food from puzzle boxes. |
| Habituation | 26 hours | Mate choice apparatus | Focal individuals were placed in a two-way mate choice apparatus. |
| Preference test phase | 4 days | Mate choice apparatus | Focal female interacted with the two focal males for 2 h each day |

#### (i) Foraging skill training phase

After being moved to the individual cages (28 × 23 × 25 cm), the focal female and both males were trained to obtain food by opening one of two types of puzzle boxes (Fig.1a): a one-step box (a petri dish where food could be accessed by opening a lid) or a two-step box (a box where food could be accessed by opening a door and pulling a drawer). Females learned the same skill as one of the two males, using a counterbalanced design for females (9 females each learned to access one of the two boxes). Each bird underwent two daily training sessions (morning: 10:00– 11:00, afternoon: 16:30–17:30), during which they could only obtain food by solving the puzzle boxes. Successful acquisition of the foraging skill was defined as the point when an individual successfully solved the task in 10 consecutive trials. All the birds successfully reached the learning criterion. Individuals were also exposed to the other puzzle box for two days after successfully learning, to ensure familiarity with both puzzle boxes, as they would interact with both types of boxes during the observation phase (see below). For males, the other puzzle box was present in the open position. For females, they could only open the puzzle box type that they had been trained on, while the other box type was sealed with tape to prevent them from acquiring additional skills. Outside the training sessions, a regular feeder (i.e., a transparent box commonly used in our lab) was available to ensure daily food intake.

#### (ii) Observation phase

The observation phase began after a two-day familiarization with both types of puzzle boxes. During the observation phase, females observed males performing their trained foraging skills on four consecutive days. Each male received both types of puzzle boxes six times during a 30-minute period between 10:00 and 11:00 AM. Thus, each male completed a total of 24 trials over the four-day long observation phase. In each trial, the puzzle boxes were presented to the males for one minute. A puzzle box of the type they had not been trained on was also present in their cage during the observation phase, but these boxes were sealed with tape to prevent the males from opening them by chance. At the beginning of the experiment, we removed one side of the opaque white barrier between the demonstrator and observer cages (Fig.1b), allowing the observing female to interact with the male. We scored the number of trials in which each male successfully opened the boxes as an indicator of male foraging performance. The subjects were food-deprived from 18:00 to the next day at 10:00.

After performing the foraging tasks, males were allowed to feed *ad libitum* from regular feeders (11:00 to 18:00). During this same period, female subjects continued their training as before to reinforce the importance of foraging skill. Females received both puzzle boxes but were only able to feed from the one they had been trained on, while the untrained box was sealed with tape.

#### (iii) Preference test

Once the observation phase was completed, the subjects were introduced into a two-way mate choice apparatus (200 × 40 × 40 cm), which consisted of a middle compartment (for the female) and two side compartments (one for each male, see Fig.2). Detailed information on the cage and procedures is given in Chen et al. (2019). Before beginning the preference tests, birds were given 26 h to habituate to the new cage, being visually isolated from each other, although not acoustically. The focal female was then allowed to choose between two males for 2 h each day (14:00–16:00) in a mate choice cage over four consecutive days. To eliminate potential side bias, the positions of males with different foraging skills were initially counterbalanced, and then they swapped sides every day. We used the time female spent in preference zones as a proxy for female preference for the respective male. After the 2-hour preference test, males had unrestricted access to regular feeders, while females continued to receive the two puzzle boxes (with the non-trained one sealed with tape) and the regular feeders, all filled with food.

**Fig. 1.**
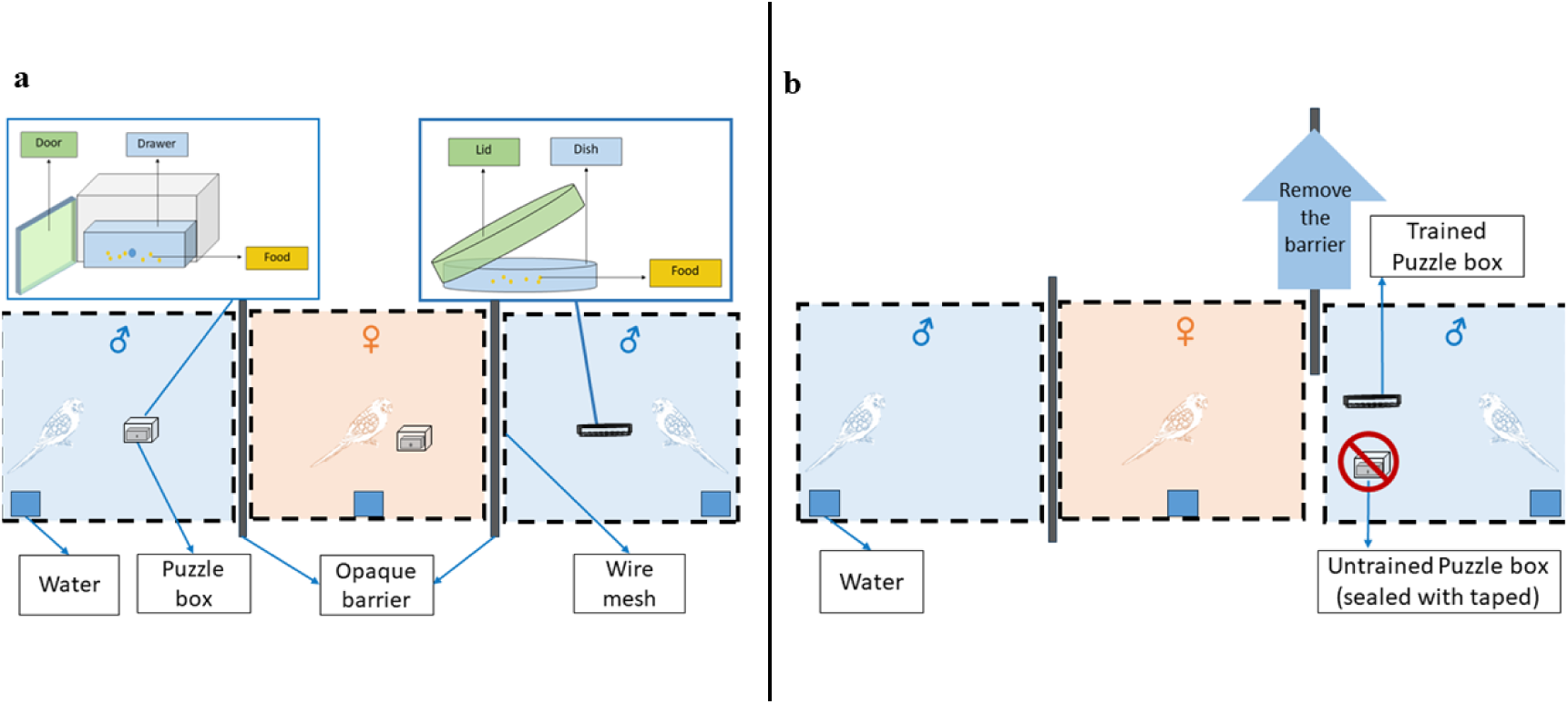
Design of the training and observing phase with foraging puzzle boxes (top view) (a) Training phase; two-step puzzle (opening a door and pulling a drawer): left blue box, one step puzzle: right blue box. (b) Observing phase; during the observing phase, one side of the opaque barriers was removed, allowing females to observe and interact with one male at a time. The puzzle boxes were only available to the male performing at that time. All cage compartments were colorless and transparent, they are here colored for illustrative purpose.

**Fig. 2.**
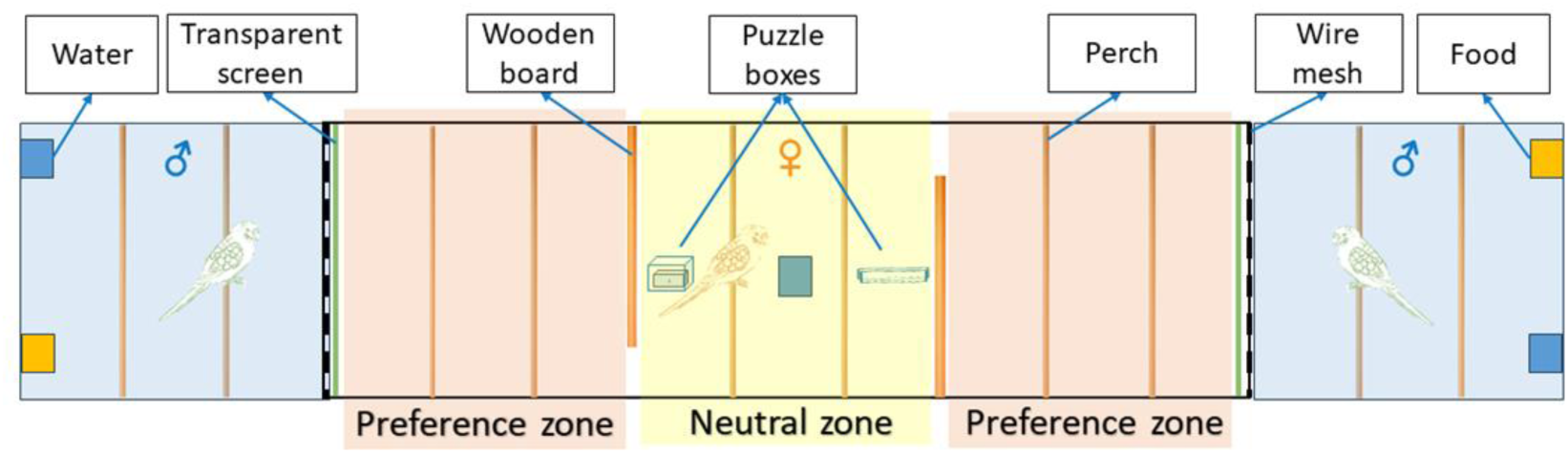
Diagram of the mate choice apparatus (top view) Preference zones and neutral zone separated by wooden boards, with entrances at different positions to prevent females from observing two males at the same time. Only females were allowed to interact with puzzle boxes to prevent their time allocation from being influenced by learning foraging skills. All cage compartments were colorless and transparent, they are here colored for illustrative purpose.

## Data analysis

Statistical analyses were performed in R software version 4.1.2 (R Core Team 2021), with a significance level set at α = 0.05. The linear mixed models (LMM) model and linear models (LM) were fit using the *lmer()* function in “lme4” package (Bates et al. 2009). The one-way ANOVA was conducted using the *aov()* function. The Principal Component Analysis (PCA) was implemented in the “nFactors” package (Ledesma and Valero-Mora 2019). The figures were drawn using GraphPad Prism 8.3.0 software (Version 8.3.0, GraphPad Software, 2019). Values are presented as means ± standard error (SE).

The time females spent in preference zones with the males was recorded as an indicator of female preference. To analyze female preferences, we used LMM to assess the effect of male foraging performance (successful number of trials of box opening during the observation phase), female skill type (one-step vs two-step box), male skill type (same vs different from females), male position in mate choice cage (left vs right) and preference test day (1^st^, 2^nd^, 3^rd^, 4^th^). Female identity was fitted as random effect. The interactions between male position and male skill type, and male position and female skill type were not significant and therefore excluded from the final model. Given that females had an overall bias for males located in the left compartments, we analyzed the data separately for the first two preference tests and the last two preference tests using similar models as described above. This approach allowed us to test whether this side bias changed across the tests. Male foraging performance, female skill type, male skill type and male position in mate choice cage were included as fixed factors, and the focal female identity was included as random effect.

We used one-way ANOVAs to test whether the latency to enter one of the preference zones for the first time differed among the four preference tests and whether the number of times females changed positions between the preference zones and the neutral zone differed across the four preference tests. To assess whether morphological characteristics of male affected female preference, we conducted a PCA to reduce the dimensionality of morphological measurements because most of them exhibited moderate to strong collinearity. The PCA with varimax rotation was conducted on the morphometric measurements of the male birds (body weight, wing length, tail length, and tarsus length) (Table. 2). Then a LM was employed to examine the influence of PC1 (body mass) and PC2 (tarsus length) on female preferences.

**Table 2.** Principal component analysis (PCA) of morphological characters in male budgerigars. Standardized loadings of the main contributors to each component are highlighted in bold.

|  | PC1 (body mass) | PC2 (tarsus length) |
| --- | --- | --- |
| Body weight (g) | **0.84** | -0.02 |
| Tarsus (mm) | 0.04 | **1.00** |
| Wing (mm) | **0.82** | 0.13 |
| Tail (mm) | **0.81** | -0.03 |
| SS loadings | 2.04 | 1.01 |
| Proportion Variance | 0.51 | 0.25 |
| Cumulative Variance | 0.51 | 0.76 |
| Proportion Explained | 0.67 | 0.33 |
| Cumulative Proportion | 0.67 | 1.00 |

## RESULTS

On average, individuals required 7.71 ± 0.28 days of training to successfully learn to open the foraging puzzle boxes. During the observation phase, males solved on average 91.05% ± 1.80 of the foraging task trials. The performance of males during the observation phase did not influence the time females spent close to the males (Table. 3). During the preference test phase, the latency of females to initially enter the preference zones (i.e., the two zones on either side of the middle compartment closest to the males) significantly decreased across the four days (*F*_1,66_ = 4.747, P = 0.033; Fig. 3a), while the number of times females changed positions between the preference zones and the neutral zone did not differ across the four days (*F*_1,66_ = 0.662, P = 0.42; Fig. 3b). Additionally, there was no significant difference in the time females spent in the neutral zone across the four preference tests (1^st^ test: 47.67 ± 8.69 min, 2^nd^ test: 34.16 ± 5.43 min, 3^rd^ test: 42.90 ± 7.57 min, 4^th^ test: 31.77 ± 6.83 min; *F*_1,66_ = 0.942, P = 0.426).

**Fig. 3.**
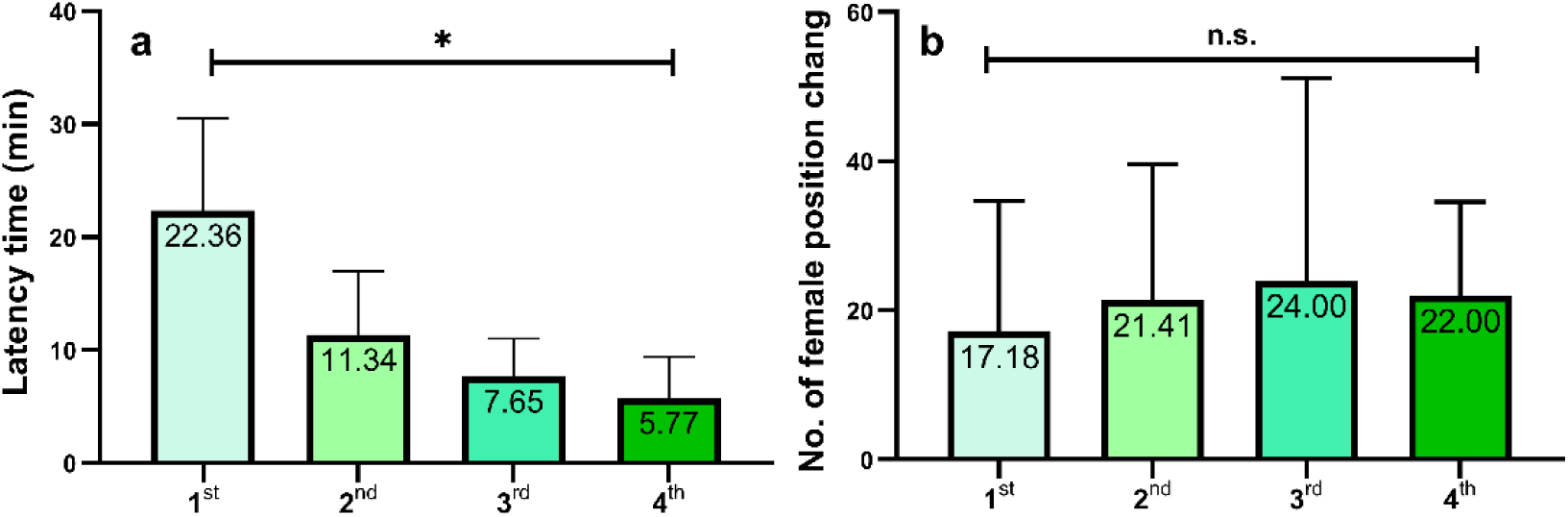
Latency to preference zone (a) and the number of times females changed positions (b) across four preference tests. The numbers displayed on the bars represent the mean values. n.s.: P > 0.05, *: P < 0.05. Bar = mean, whisker = standard error.

Overall, females spent more time close to males that had a different foraging skill than they had (Table 3; Fig. 4a), but 13 out of 17 of females had a side bias, and preferred males located in the left compartment of the apparatus (Table 3; Fig. 4b). Given that the location of the males (left vs right compartment) was changed every day, we compared female preference during the first two day vs. the subsequent two days to account for this side bias. During the first two days, only male position was associated with female preference (LMM: male position: ﹣28.23 ± 8.52, χ^2^ =10.98, P < 0.001; male skill: 7.43 ± 8.66, χ^2^ =0.74, P = 0.39; Fig. 5). However, during the subsequent two days, only male skill type was associated with female preference (LMM: male position: ﹣14.72 ± 8.93, χ^2^ = 2.72, P = 0.10; male skill: ﹣18.56 ± 9.08, χ^2^ =4.18, P = 0.041; Fig. 5).

**Fig. 4.**
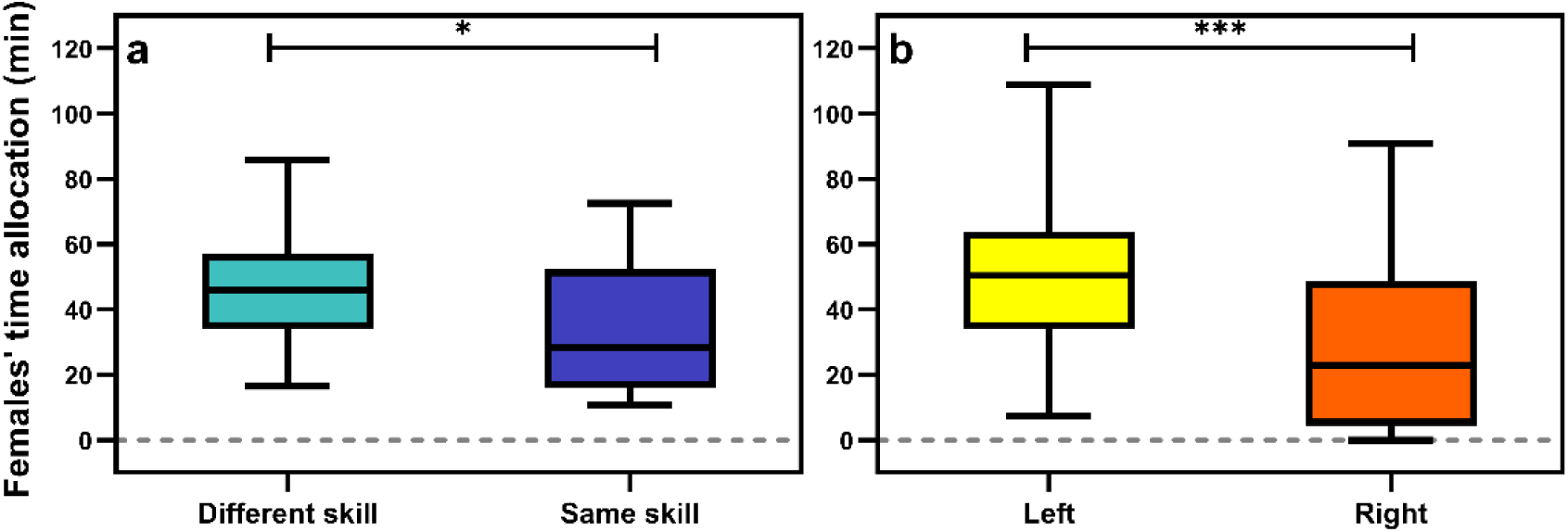
Females’ time allocation to males with same and different foraging skills and left/ right preference zones. **(a)**: Time allocated by focal female budgerigars (N=17 female × 4 tests) to different-skill males (47.25 ± 4.33 min; green) and same-skill males33.62 ± 4.83 min; purple). **(b)**: to left side (51.18 ± 6.74 min; yellow) and right side (29.70 ± 6.84 min; orange). The figure shows the time allocation of females to different-skill males in green and to same-skill males in purple. *: P < 0.05, **: P < 0.01, ***: P < 0.001. Line = median, box = 25th to the 75th percentiles, whisker = min & max.

**Fig. 5.**
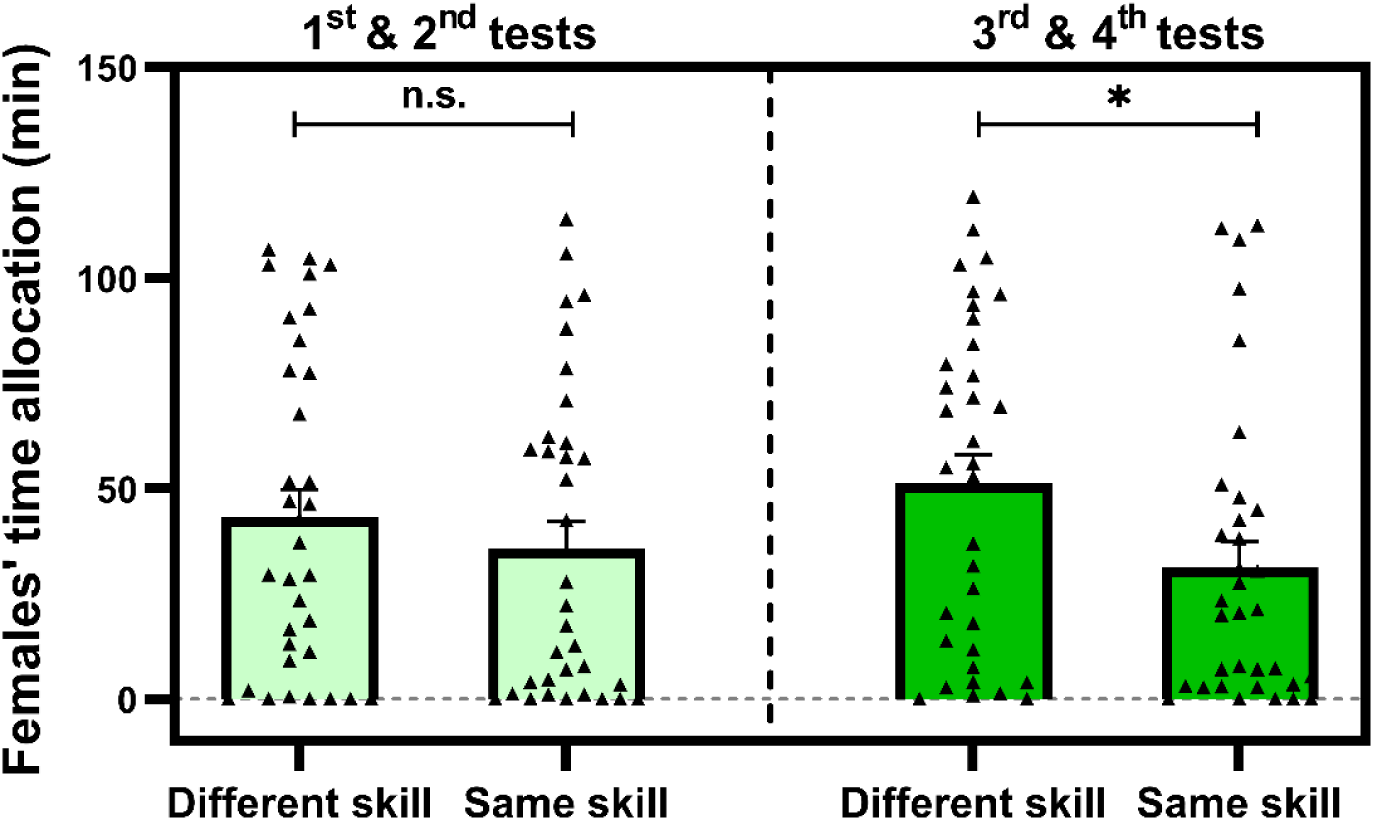
**Females’ time allocation to males with same and different foraging skills in fist two tests (light green) and the last two tests (dark green) n.s.**: P > 0.05, *: P < 0.05. Raw values are indicated with black triangles. Bar = mean, whisker = standard error. Female preference was not influenced by puzzle box type (one step box males: 38.28 ± 4.68, two step box males: 42.59 ± 4.49; paired t-test: t = 0.67, P = 0.51; Fig. 6). Likewise, female preferences were not affected by their own foraging skill types (Table 3), and neither did morphological characteristics of males influence female preference (LM: PC1 (body mass): -10.68 ± 13.40, t = -0.80, P = 0.43; PC2 (tarsus length): -25.25 ± 13.40, t = -1.88, P = 0.07).

**Table 3.** LMM analysis assessing the effect of female skill types, male skill types, male positions, male performance and preference tests on female preference (i.e., the time female allocated to males). Female ID was included as random effect. Significant parameters are highlighted in bold.

| Parameter | Estimate | SE | $\chi^2$ | P |
| --- | --- | --- | --- | --- |
| **Preference test times (s, n = 136)** |  |  |  |  |
| Intercept | 19.22 | 28.23 |  |  |
| Female skill<br>(two-step box vs petri dish) | 9.16 | 6.25 | 2.15 | 0.14 |
| **Male skill**<br>**(same vs different skill)** | **13.00** | **6.23** | **4.35** | **0.037\*** |
| **Male position**<br>**(left vs right)** | **- 21.48** | **6.13** | **12.27** | **< 0.001\*\*\*** |
| Male performance | 17.28 | 30.58 | 0.32 | 0.57 |
| Preference tests (1<sup>st</sup>, 2<sup>nd</sup>, 3<sup>rd</sup>, 4<sup>th</sup>) | 1.95 | 2.74 | 0.51 | 0.48 |

## DISCUSSION

Our results show that female budgerigars prefer males with foraging skills that differ from their own. However, the type of foraging skill *per se* (Fig.6; one-step vs two-step box) and the number of times males successfully solved foraging puzzle boxes during the observation phase did not influence female preference. Across the four days of the preference test, females showed a notable positional bias during the first two days, but a preference for males with different foraging skills in the subsequent two days (Fig. 5). Moreover, while the latency to enter the preference zone decreased across the preference tests, the time spent in the neutral zone did not change. These results suggest that females repeatedly evaluated the males, allowing them to overcome the initial positional bias to prefer males with different foraging skills.

**Fig. 6.**
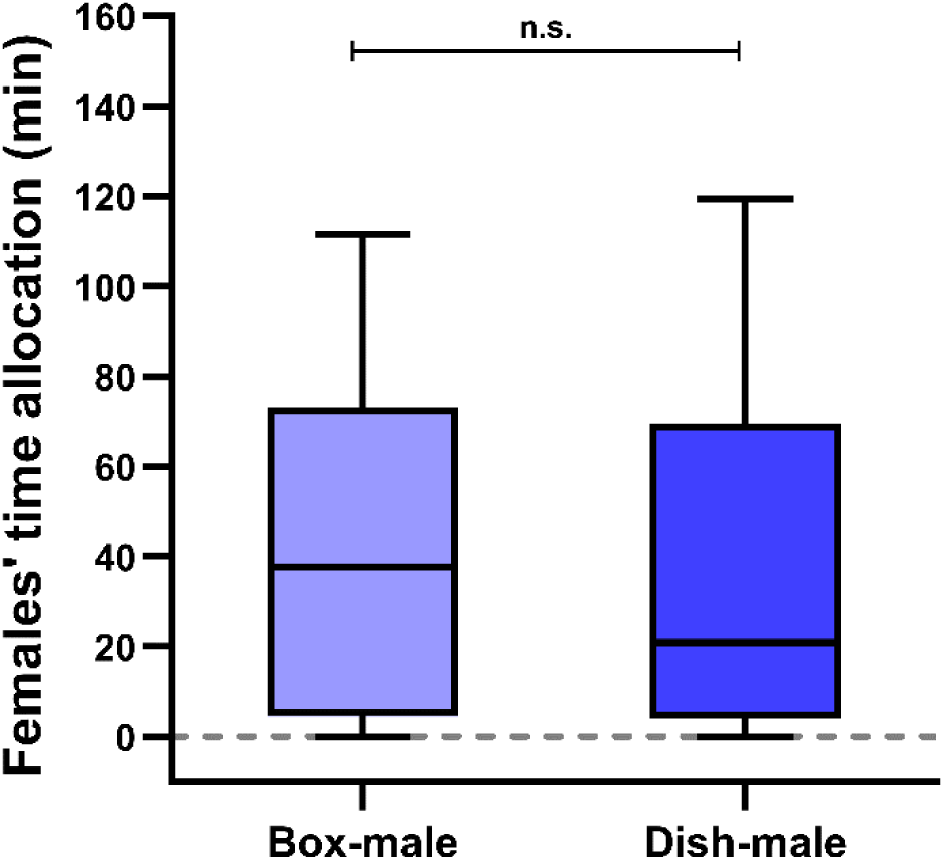
Time spent by female budgerigars with males exhibiting different foraging skills. The time females (N=17× 4 tests) spent with the males that could open the petri dish (Dish-male: 38.28 ± 4.68 min) and the males that could open the two-step box (Box-male: 42.59 ± 4.49 min). Line = median, box = 25th to the 75th percentiles, whisker = min & max.

The skill-pool hypothesis posits that individuals in group living species should join groups of conspecifics that differ in their foraging behaviors (Giraldeau 1984). Previous studies have shown that individuals indeed benefit from foraging with others who can access different food sources (Aljadeff et al. 2020; Morand-Ferron and Quinn 2011). Our study provides the first support for this hypothesis in mate choice context, by showing a disassortative mating pattern related to foraging skills. Choosing mates with different foraging skills can be beneficial for two reasons. First, young learn their foraging skills socially from caregivers particularly in species where offspring remain at least some time with their parent(s) beyond nutritional independence (Carvajal and Schuppli 2022; Uomini et al. 2020). For example, a cross-fostering experiment on social learning of foraging in blue tits (*Cyanistes caeruleus*) and great tits (*Parus major*) revealed that early learning from foster parents caused a shift in foraging techniques towards the foster species, lasting for life and even influencing subsequent generations (Slagsvold and Wiebe 2011). Thus, in species with biparental care, pairing with different-skilled individuals could contribute to the social transmission of a more diverse foraging skillset from parents to their offspring (van Schaik 2010). Second, choosing a mate with different foraging skills can increase within-pair foraging success via foraging niche diversification and reduced competition over food. For example, body size differences between females and males occur in many raptors and owls, which has been suggested to reduce within-pair competition over prey (Krüger 2005). Similarly, within-pair differences in foraging skills likely broaden the foraging niche, and allow parents to ensure adequate offspring provisioning. A study on zebra finches (*Taeniopygia guttata*) demonstrated that paired males tended to avoid the food their mate selected, indicating a strategy to reduce competition with their partner (Templeton et al. 2017). Thus, disassortative mate choice with respect to foraging skills can broaden the foraging skill set of offspring, which might contribute to fitness (Boogert et al. 2011).

Our findings contrast with a number of studies that show assortative patterns of foraging skills. Experimental studies showed that individuals adopt local foraging norms, as observed in vervet monkeys (*Chlorocebus aethiops*) (van de Waal et al. 2013) and great tits (Aplin et al. 2014). We note that both of these studies assessed the avoidance of a specific food type, respectively the learning of a specific skill outside the mate choice context, rather than foraging skillset diversity. Thus, further studies are required to assess the factors associated with disassortative and assortative mate choice patterns with respect to foraging skills (Camacho-Alpízar et al. 2020).

Our experiment assessed mate preference, but did not assess the subsequent mate choice decision. We note that preferences for other individuals can also reflect social preferences outside the mating context (e.g., Engeszer et al. 2004; Seyfarth and Cheney 2012). However, a previous study on budgerigars demonstrated that focal females shifted their preference to initially less-preferred males after those males acquired foraging skills, as indicated by the increased time females spent in proximity to them (Chen et al. 2019). In contrast, focal females did not alter their preference for initially less-preferred females, when females acquired similar foraging skills as these males. This suggests that the acquisition of foraging skills could influence female mate preferences rather than social preferences.

It is established that kinship and social familiarity can influence social preferences (e.g., Griesser et al. 2015). To exclude a role of these factors, we sourced experimental females and males from different suppliers to ensure they were neither related nor familiar with each other. Also, all individuals in our experiment were unpaired. Accordingly, it seems reasonable to assume that the social preference of females for males with different foraging skills also reflected mate choice preferences, although other factors could also contribute to this behavior. Budgerigars typically remain with their mates throughout the year and even when flocking, they are usually in close proximity to their mates and synchronize their movements (Fritz 1976). Moreover, the mate choice apparatus used here is widely used to study mate choice in birds (including budgerigars; e.g., Arnold et al. 2002; Zampiga et al. 2004). Thus, the preference of female budgerigars for male with different foraging skills is likely related to mating behavior.

In this study, female budgerigars exhibited a side bias in the mate choice apparatus, which may be attributed to the natural lateralization of spatial attention. Birds are known to display a left-side bias in spatial tasks, likely due to the dominance of the right hemisphere in controlling spatial attention (Diekamp et al. 2005). In budgerigars, individual behavioral lateralization has been observed in various tasks (Schiffner and Srinivasan 2013). Additionally, asymmetries in the experimental room and lighting conditions could further enhance this side-bias. However, it is important to note that our setup controlled for this side bias, and that further studies are required to understand the reasons underlying this side bias *per se*.

In conclusion, our study reveals that i) similarity in foraging skills influences social preference for mates and therefore likely also mate choice; ii) females considered the foraging skills of the males in relation to their own foraging skills; iii) females repeatedly assessed the males’ characteristics over time; and iv) female preferences were not influenced by the types of foraging puzzle boxes. Our findings suggest that budgerigars can associate specific foraging skills with particular conspecifics. Accordingly, female mate preferences in budgerigars can broaden the foraging skill repertoire of pairs, facilitating the exploitation of additional food resources, and thereby supporting the evolution of foraging skill diversity. Alternatively, If the observed pattern would reflect female social preference, it could still broaden their foraging niche of a pair, making it more likely for their offspring to acquire new foraging skills.

## Funding

This study was financially supported by the Natural Science Foundation of China (No. 31501868). MG was supported by a Heisenberg Grant GR 4650/2-1 by the German Research Foundation DFG.

## Ethical approval

The study was approved by the Animal Care and Use Committee of the Institute of Zoology, Chinese Academy of Sciences (CAS). All experimental procedures complied with applicable governmental regulations concerning the ethical use of animals and biodiversity rights.

## Acknowledgments

We thank Ying-qiang Lou for helping with the experiments, Richard Gunner for improving the English. We are grateful to Carel van Schaik for his comments on the manuscript. We also thank two anonymous reviewers and the editor for their valuable comments and suggestions.

## Notes

### Competing Interest Statement

The authors have declared no competing interest.

